# Humanization of the *rpb9* locus in fission yeast reveals conserved and divergent roles of *rpb9* and human *POLR2I*

**DOI:** 10.64898/2026.04.02.716003

**Authors:** Jared M. Finkel, Micah G. Williams, Mamta B. Nirmal, Samakshi Pandey, Erik D. Howe, Cameron T. Liu, Jeremy R. Lohman, Nimisha Sharma, Tommy V. Vo

## Abstract

**Background/Objectives:** RNA polymerase II is a multifunctional complex that is critical for gene regulation and environmental responses. Its POLR2I subunit in human is associated with various pathologies, including cancer chemoresistance. However, much of our understanding of how POLR2I could function indirectly derives from studies of its homologs in yeasts called Rpb9. Here, we endogenously humanized the *rpb9* gene of the fission yeast *Schizosaccharomyces pombe* to examine the functional capabilities of POLR2I.

**Methods:** We edited the genomic *rpb9* locus in *S. pombe* so that it encodes the human POLR2I protein, and investigated functional and structural conservation.

**Results:** With our humanized yeast system, we find widespread functional complementation by human *POLR2I* of *S. pombe rpb9* roles in yeast growth, chronological aging, and stress responses. We also find that *POLR2I* complements novel roles for yeast *rpb9* in facultative heterochromatin assembly, resistance against the chemotherapy 5-fluorouracil, and resistance against the fungicide thiabendazole. In contrast, we find that *POLR2I* cannot complement the role of *rpb9* in resistance against the transcription elongation inhibitor 6-azauracil (6-AU) in our system. Interestingly, *POLR2I* could complement 6-AU resistance if ectopically expressed. Lastly, we observe extensive structural homology between Rpb9 and POLR2I proteins.

**Conclusions:** Our study establishes an endogenous cross-species gene complementation strategy that uncovers both conserved and rewired functions of fission yeast *rpb9* and its human homolog, *POLR2I*. In addition to validating conserved roles, we also identified conservation of previously unrecognized roles of *rpb9* in heterochromatin formation and chemoresistance.

## 1. Introduction

Gene regulation is a fundamental process that shapes cell fate and behaviors. In eukaryotes, gene expression is prominently regulated at the level of chromatin through transcriptional [1], co-transcriptional [2], and post-transcriptional mechanisms [3]. DNA-dependent RNA polymerase II (Pol II) plays a central role in these processes by functioning both as an enzyme and as a molecular recruitment platform [4]. As an enzyme, Pol II mediates transcription by synthesizing RNA [5–8]. As a platform, it can accommodate diverse post-translational modifications and interact with other biomolecules to coordinate transcription, chromatin modifications, chromatin remodeling, and RNA processing [9–11].

Pol II comprises 12 core protein subunits that collectively mediate its distinct functions [12–15]. While most Pol II subunits are essential for viability in yeast and metazoans, the Rpb9 subunit (also known as POLR2I in humans) is non-essential but is involved in environmental responses [16,17]. Most of our understanding of the molecular functions of Rpb9 comes from studies in the model budding yeast *Saccharomyces cerevisiae*, which indicate that Rpb9 plays important roles in transcription start site selection, transcription elongation, transcriptional fidelity, and transcription-coupled DNA repair [18–21]. Rpb9 performs these roles, in part, by modulating interactions of Pol II with transcription factors such as TFIIS [20] and TFIIF [22]. In contrast, there is a paucity of studies that have directly interrogated the molecular functions of human POLR2I [23]. This limits our ability to directly translate knowledge of yeast Rpb9 to its human homolog POLR2I. As recent studies showed that POLR2I amplification or upregulation is associated with colorectal cancer metastasis [24], chemoresistance of head and neck cancer [25], and hypertensive nephropathy [26], elucidating its molecular roles could advance our understanding of the etiology of these conditions.

Cross-species gene complementation is a powerful genetic approach for assessing the functional conservation of homologous coding genes [27–30]. For a given gene pair in which one gene encodes a protein with established function(s) and the other encodes a homolog with unclear function(s), the better-characterized gene is replaced with the less-characterized counterpart [31]. Functional assays are then used to determine which known activities can be complemented by the uncharacterized homolog. Successful complementation provides strong genetic evidence that the two genes (likely via their encoded proteins) share the function(s) in question. Gene complementation studies have been conducted to test whether human *POLR2I* can complement the roles of *rpb9* in the model yeasts *S. cerevisiae* and *Schizosaccharomyces pombe*. However, the biological interpretation of some results has been challenging. For example, high-level expression of *POLR2I* suppressed the hypersensitivity of yeast lacking *rpb9* (*rpb9Δ*) to elevated temperature, whereas lower expression levels produced no complementation [17,32]. This makes it unclear whether yeast *rpb9* and human *POLR2I* genuinely share temperature-related roles or if the complementation is rather context-dependent. Such challenges stem from the design of previous gene complementation studies [31,33], which relied on ectopic expression of *POLR2I* from plasmids in *rpb9*Δ yeast cells, followed by phenotypic analysis [17,34]. Although this strategy has enabled powerful functional interrogation of human genes in genetically tractable yeast systems, it also introduces confounding variables, including disruption of native regulatory context, non-physiological expression levels (determined by plasmid copy number or artificial promoters), and dependence on selective pressures to maintain plasmids. Refining the design of gene complementation studies for *rpb9^POLR2I^* to minimize these extraneous variables would strengthen the biological interpretation of their functional relationships.

In this study, we developed a genomic gene complementation strategy to functionally re-examine human *POLR2I* in the fission yeast *Schizosaccharomyces pombe*. This approach overcomes limitations of traditional plasmid-based systems by directly replacing the endogenous *rpb9* gene with *POLR2I*, at the native *rpb9* locus, thereby preserving the native regulatory context of the *rpb9* locus. Genomic integration also enables analysis of *POLR2I* function under standard non-selective rich-media conditions. Consistent with previous plasmid-based complementation studies [17,34], we find that endogenously expressed *POLR2I* robustly complements *S. pombe rpb9* roles in cell growth in the presence of several stress conditions. In addition, we identify a previously unrecognized and conserved role for *rpb9^POLR2I^* in the formation of facultative heterochromatin in *S. pombe*. Notably, our system reveals a lack of complementation in response to 6-azauracil (6-AU), a transcription elongation inhibitor, suggesting partial functional divergence between fission yeast *rpb9* and human *POLR2I*. Collectively, our study establishes a physiologically relevant gene complementation framework that both reproduces and extends current understanding of *POLR2I* function.

## 2. Materials and Methods

### 2.1. Yeast culturing and manipulation

*S. pombe* yeast strains that were used in this study are listed in Table S1. The *rpb9*Δ yeast strain was constructed by replacing the *rpb9* open reading frame with the *kanMX* gene by using a PCR-based gene deletion procedure with transformation by lithium acetate method, as previously described [35]. The *kanMX* gene was derived from a pFA6a- *kanMX* plasmid that was purchased from Addgene (Addgene 39296). All yeast were cultured in media that were based on the yeast extract rich medium with glucose and adenine supplement (YEA; 30 g/L glucose, 5 g/L yeast extract, 75 mg/L adenine, pH 5.5) and incubated at 32°C. To make plates, YEA-based media was prepared with 2% agar. Liquid cultures were grown at 32°C with 220 rpm shaking for aeration. For our experiments, we have used YEA media with the following: 5-FOA (GoldBio, cat no. F-230, 850 mg/L), G418 sulfate (GoldBio, cat no. G-418, 50 mg/L), NaCl (Sigma, cat no. 746398), 5-fluorouracil (Sigma, cat no. F6627, 25 μM), thiabendazole (Fisher Scientific, cat no. AAJ6000909, 20 μg/mL), and 6-azauracil (Fisher Scientific). For experiments using cells that carried exogenous expression plasmids, cell manipulation and spotting assays were performed as previously described in [17].

### 2.2 Endogenous humanization of rpb9 in S. pombe

To integrate either *POLR2I* or *rpb9* into the endogenous *rpb9* locus, we started with a wild-type yeast strain that has an intact *rpb9+* locus, and lacks the *ura4* and *kanMX* genes. First, we constructed a pFA6a-*ura4*-*kanMX* plasmid by cloning the *ura4* gene, along with 500 bp upstream and downstream regions, into the commercial pFA6a-*kanMX* plasmid (Addgene 39296). The upstream and downstream regions include the native ura4 promoter and termination sequences [36]. PCR amplification of the *ura4* locus was performed to add AscI and BglII restriction enzyme sites onto the PCR amplicon ends. Subsequently, conventional restriction enzyme cloning was performed to integrate the *ura4* amplicon into pFA6a-*kanMX*, placing it upstream of the *kanMX* promoter. The resultant pFA6a-*ura4*-*kanMX* plasmid was verified by Sanger sequencing.

Next, PCR-based homologous recombination [35] was used to replace the endogenous *rpb9* open reading frame DNA sequence with *ura4*-*kanMX*. The *ura4-kanMX* cassette was PCR amplified using Splicing by Overlapping Extension (SOE) PCR [37] to append ∼300 bp of sequence homology to the genomic region that flanks the *rpb9* open reading frame. Cells, where the swap was successful, grew on Pombe Minimum Glutamate (PMG) media lacking uracil and on YEA media with G418, but failed to grow on YEA media with 5-FOA.

Finally, PCR-based homologous recombination [35] was used to replace the *ura4-kanMX* cassette with the open reading frame DNA sequence of *POLR2I* or *rpb9*. The *POLR2I* DNA sequence was codon-optimized for *S. pombe* and synthesized as a double-stranded DNA product by the Integrated DNA Technologies (IDT) company. The *rpb9* DNA sequence was PCR amplified from the ancestral wild-type strain TVTV39. Cells that successfully lost the *ura4-kanMX* cassette grew on YEA media with 5-FOA, but failed to grow on PMG media lacking uracil and YEA media with G418. The genotypes of the resultant yeast strains were verified by PCR and Sanger sequencing analyses. The DNA sequence of the POLR2I open reading frame used in this study is provided in Table S2.

### 2.3. Yeast spotting assays

Yeast cells were pre-cultured at 32°C for 220 rpm overnight until cells reached log growth phase. Afterwards, cell optical densities (OD_600_) were measured using a Denovix DS-11+ apparatus and then cells were normalized to the equivalent OD_600_ of 0.2 – 0.5. Finally, cells were serially diluted 4-fold or 10-fold, and equal volumes of cells were spotted onto the indicated media plates. Spotted plates were incubated at 32°C without shaking for 2-5 days. Images were collected using a BioRad Chemidoc imager using the Colorimeteric setting. Any image correction was uniformly applied to the entirety of the shown images.

### 2.4. Western blotting

Western blotting was essentially performed as previously described [38]. Specifically, approximately two OD_600_ units of yeast cells were harvested by centrifugation into 2 mL screw-cap tubes. The cell pellet was resuspended in 200 µL of cold 20% trichloroacetic acid (TCA). Approximately 400 µL of acid-washed glass beads were added, and the cells were lysed using mechanical disruption with a bead beater (20 seconds beating × 3 cycles with 45-second intervals on ice). Following lysis, each tube was punctured, placed in a 5ml polystyrene collection tube, and the lysate was collected by centrifugation at 1000 rpm for 30 seconds. The remaining beads were washed with 400 µL of cold 5% TCA, spun again, and the wash was pooled with the initial lysate. The combined lysate (∼600 µL total) was transferred to a 1.5 mL microcentrifuge tube and centrifuged at 13,000 rpm for 5 minutes at 4°C to pellet the proteins. Supernatant was discarded and centrifuged 2-3 times to remove residual TCA. The resulting protein pellet was resuspended in 100 µL of SDS-PAGE 1x sample loading buffer (Thermo Fisher Scientific, cat no. NP0007-2988243). Pellets were often difficult to dissolve and required heating to fully solubilize. Samples were then boiled at 95°C and loaded on SDS-PAGE 4–12% Bis-Tris gels (Thermo Fisher Scientific, cat no. 25010870). Electrophoresis was performed in an Invitrogen mini tank apparatus using NuPage 1X running buffer at room temperature at 200 volts for 45 minutes. Proteins were transferred onto PVDF membranes (pre-activated with methanol) using wet transfer (Nupage 1X transfer buffer (NP0006-2988243) at room temperature.

Following the transfer, membranes were stained with Ponceau S stain solution (Thermo Fisher Scientific, cat no. A40000279) to reveal total protein. A portion of the staining pattern was used as a measure of the loading control. Then, the stain was washed away and then membranes were blocked for 1 hour at room temperature in 1X TBST (Tris-buffered saline with 0.1% Tween-20) containing 5% non-fat dry milk. Membranes were then incubated overnight at 4°C with 1:2,000 mouse monoclonal anti-GFP (Sigma, cat no. 11814460001, clones 7.1 and 13.1) that was diluted in blocking buffer. After washing three times with 1X TBST (10 minutes each), membranes were incubated with an HRP-conjugated secondary anti-mouse (Jackson laboratory, cat no. 115-035-146, at 1:5,000 dilution) for 1 hour at room temperature. Following three additional 1X TBST washes, signal detection was performed using an enhanced chemiluminescence (Cytiva, cat no. RPN2232-18206643) substrate and visualized using the BioRad chemiluminescence imaging system.

### 2.5. Live-cell microscopy

To visualize Rpb9 or POLR2I localization in yeast cells, we used strains where the *gfp* gene was genomically integrated adjacent to the *rpb9* or *POLR2I* open reading frames to encode for C-terminally tagged Rpb9-GFP or POLR2I-GFP fusion proteins. Yeast cells were grown in YEA media to the exponential growth phase and then 1 OD_600_ of cells were washed with phosphate-buffered saline (PBS) by pelleting cells in a centrifuge at ≥13,000g for 1 minute, for three times. To visualize nuclear DNA, cells were stained with Hoescht 33342 by adding the dye to create a 1µg/mL solution. Cells were then incubated in darkness for 15 minutes. Next, cells were added to a glass slide and visualized using the Brightfield, DAPI, or eGFP fluorescence settings of the ZEISS Axio Observer Z1 inverted fluorescence microscope using the 63x oil immersion objective. Images were processed using the ZEN Pro software (ZEISS) software.

### 2.6. Chronological aging assay

Yeast cells were cultured in YEA liquid media at a constant 1:5 volume-to-flask maximum volume ratio, at 32°C with 220 rpm shaking for aeration. To assess chronological lifespan of the various yeast strains, cultures were left shaking continuously without changing the media. Day-0 is defined as the day when cultures were in exponential growth phase. Day-1 is defined as the start of the stationary phase. Day-1+ is defined as the days after initially reaching the stationary phase. At various days, cells were normalized by OD_600_ across strains, then serially diluted 1:4, and finally spotted onto YEA media plates. Plates were incubated at 32°C for 3 days to assess strain viability based on yeast growth patterns.

### 2.7. Chromatin immunoprecipitation

Chromatin immunoprecipitation was performed as previously described [38]. Specifically, *S. pombe* cells were inoculated in 50 mL of YEA medium at 32°C with shaking at 220 rpm to an OD600 of 0.5–0.6. Cells were crosslinked with 1% formaldehyde for 20 minutes at room temperature, followed by quenching with 2.5M glycine. Next, cells were washed twice with 20 mL of cold 1× PBS. The cell pellet was resuspended in 350 µL of ChIP lysis buffer (50 mM HEPES pH 7.5, 140 mM NaCl, 1 mM EDTA, 1% Triton X-100, 0.1% sodium deoxycholate, EDTA free 1X cOmplete) and disrupted with 1 mL of glass beads using bead-beating (1 min beating, 2 min on ice, repeated 3 times). Chromatin was sheared using a QSonica Q800R sonicator (20 sec ON / 40 sec OFF cycles, 70% amplitude) for 10 minutes at 4°C. Cell debris was removed by centrifugation at 4°C, 1,500×g for 5 minutes. The supernatant was transferred to fresh 1.5 mL tubes and diluted with ChIP lysis buffer. Lysates were precleared with 20 µL of protein A/G agarose beads (Santa Cruz, cat no. A3124) and incubated at 4°C for 1 hour. 50 µL of pre-cleared lysate was reserved as the 5% input control. The remaining lysate was incubated with the H3K9me2 (Abcam, cat no. ab115159) antibody and protein A/G agarose beads overnight at 4°C. Beads were washed sequentially (twice each) with the following buffers: ChIP Buffer I (50 mM HEPES pH 7.5, 140 mM NaCl, 1 mM EDTA, 1% Triton X-100, 0.1% deoxycholate), ChIP Buffer II (same as Buffer I, but with 500 mM NaCl), ChIP Buffer III (10 mM Tris-HCl pH 8.0, 250 mM LiCl, 0.5% NP-40, 0.5% deoxycholate, 1 mM EDTA), and 1x Tris-EDTA buffer.

Immunoprecipitated chromatin was extracted by incubating the agarose beads with 100 µL of elution buffer (50 mM Tris-HCl pH 8.0, 10 mM EDTA) at 65 °C and agitating at 1,000 rpm for 30 minutes. This step was repeated to obtain a total volume of 200 µL eluant. To each ChIP sample, 4 µL of 5N NaCl and 1µL RNase A (Thermo Fisher Scientific, 20 mg/mL) was added, and samples were incubated overnight at 65°C in a water bath to reverse crosslinks. Input samples were adjusted to 200 µL with elution buffer, followed by the addition of 2.6 µL of 5N NaCl and 1 µL RNase A, and incubated overnight at 65°C in a water bath to reverse crosslinks. After the sample tubes were cooled to room temperature, 1µL of Proteinase K (Thermo Fisher Scientific, 20 mg/mL) was added to each tube and incubated at 50°C for 1 hour. DNA was purified by ethanol precipitation using 3 M sodium acetate, 1 µL glycoblue (Thermo Fisher Scientific, 15 mg/mL) and resuspended into 50 µL of molecular-grade water. For quantitative PCR (qPCR), 1 µL of ChIP DNA was used per reaction with Power SYBR Green PCR master mix (Life Technologies, cat no. 4367659). Oligos used for PCRs are listed in Table S3.

### 2.8. Sequence alignment analyses

Gene sequences for *S. pombe rpb9* and human *POLR2I* were collected from Pombase and UCSC Genome Browser, respectively. Protein sequences for *S. pombe* Rpb9 were found from Pombase and human POLR2I was found from Uniprot. Visualization of gene sequence comparison and amino acid sequence comparison was generated using the Needleman-wunch algorithm from EMBOSS NEEDLE Pairwise Sequence Alignment (PSA). Values of sequence comparison for gene sequence and amino acid sequence were collected using the Clustal Omega tool from EMBL-EBI.

### 2.9. Protein structure analyses

Empirical protein structures for *S. pombe* Rpb1, Rpb2, and Rpb9 were downloaded from structure 3H0G-A, 3H0G-B, and 3H0G-I from the Protein Data Bank (PDB) database, respectively. The human POLR2I empirical structure was downloaded from 9EHZ-F from the PDB. AlphaFold predicted structures of *S. pombe* Rpb9 and human POLR2I were downloaded from the AlphaFold database AF-O74635-F1 and AF-P36954-F1, respectively [39]. RMSD, TM-Score, and visualizations were determined by comparing structures using the Pairwise Structure Alignment tool from the PDB or PyMOL. To model the Rpb1-Rpb2-Rpb9 and Rpb1-Rpb2-POLR2I complexes, AlphaFold server was used with the following sequences; *S. pombe* Rpb1 (UniProt P36594) truncated to reside 1555, *S. pombe* Rpb2 (Q02061), and *S. pombe* Rpb9 (O74635) or human POLR2I (P36954). Three zinc ions were also included in the Rpb9 and POLR2I models.

## 3. Results

### 3.1. Swapping the endogenous rpb9 gene for PORL2I in the S. pombe genome

To humanize *rpb9* in *S. pombe* cells, we performed a two-step, scarless genome swapping procedure to replace the entire *rpb9* open reading frame (ORF) with an ORF corresponding to human *POLR2I* (**Figure 1A**). First, PCR-based homologous recombination was used to swap out the *rpb9* ORF for a *ura4-kanMX* DNA cassette. Then, another round of PCR-based homologous recombination was used to replace the *ura4-kanMX* cassette with the *POLR2I* ORF to generate the humanized yeast strain (*rpb9*Δ*::POLR2I*) (Table S2). The intermediatory *ura4* and *kanMX* genes permitted selection and counter-selection of our yeast constructs (**Table 1**).

**Figure 1.**
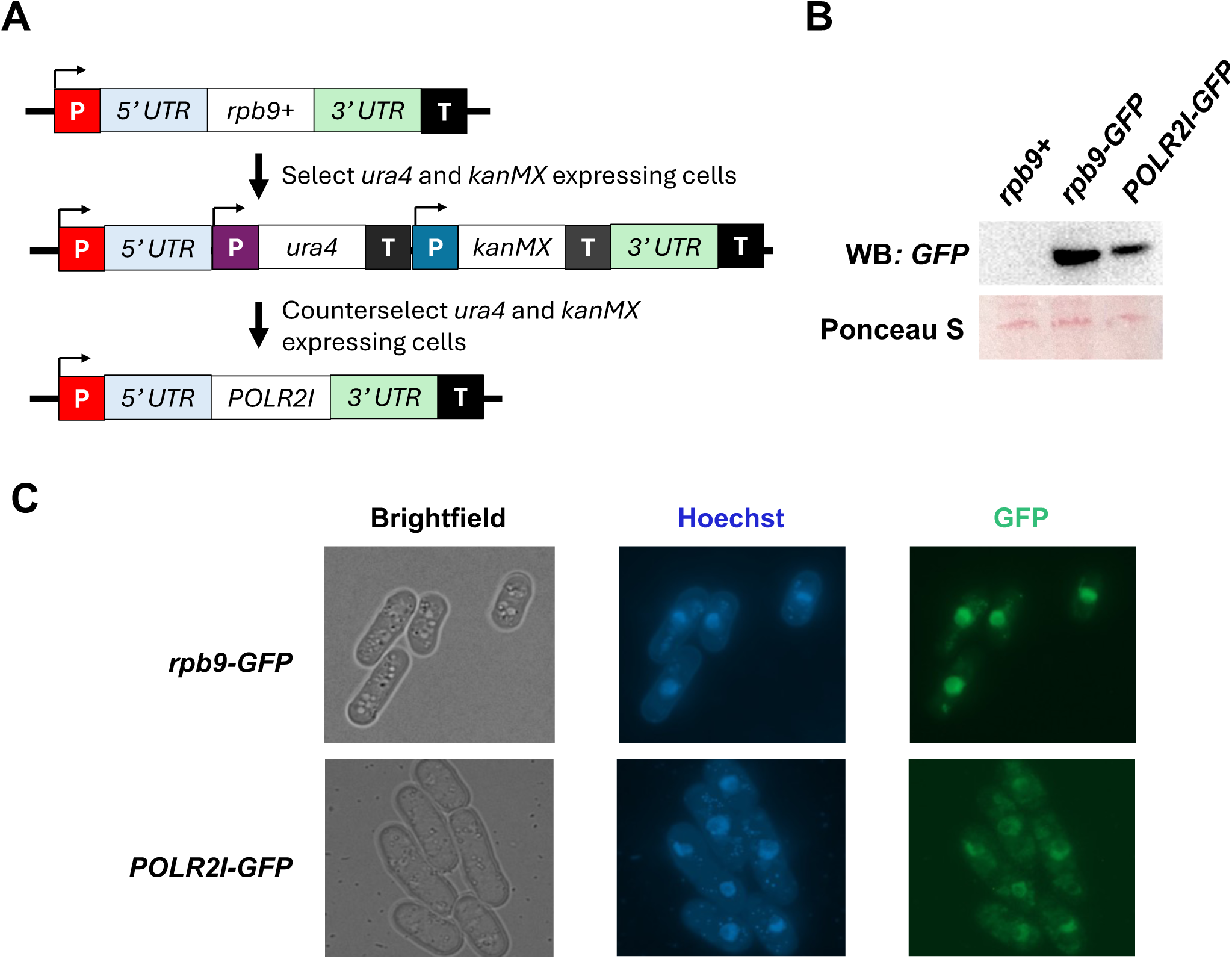
Genome-integrated *POLR2I* expresses nuclear-localized protein in *S. pombe*. (**A**) Illustration of our procedure to endogenously swap yeast *rpb9* for human *POLR2I*. (**B**) Western blot analysis of GFP tagged endogenous Rpb9 or POLR2I protein in *S. pombe* cells. Wild-type cells without GFP tag was used as a negative control. (**C**) Brightfield and fluorescence microscopy of *S. pombe* cells with GFP-tagged Rpb9 or POLR2I protein. Hoechst stains nuclear DNA.

**Table 1.** *S. pombe* growth phenotypes during our procedure to endogenously replace *rpb9* for *POLR2I*.

| Genotype | Expected growth in the<br>presence of G418 drug<sup>1</sup> | Expected growth in the<br>presence of 5-FOA drug<sup>2</sup> |
| --- | --- | --- |
| *rpb9+* | Not viable | Viable |
| *rpb9Δ::ura4-kanMX* | Viable | Not Viable |
| *rpb9Δ::POLR2I* | Not viable | Viable |

<sup>1</sup> *S. pombe* cells expressing the *kanMX* gene are resistant to the G418 antibiotic [40]. 221
<sup>2</sup> Yeast cells expressing the *ura4* gene are sensitive to compound 5-fluoroorotic acid (5-FOA) [38,41]. 222

Throughout this procedure, PCR was used to confirm successful completion of the two steps (**Figure S1**). We also validated the integration of *POLR2I* by Sanger sequencing. The humanized *rpb9*Δ*::POLR2I* yeast strain preserves the endogenous promoter, 5’- and 3’- untranslated regions (UTRs) at the original *rpb9* genomic locus. Effectively, we constructed a truly humanized *S. pombe* strain that carried a *POLR2I* ORF sequence in place of the *rpb9* ORF sequence, within the yeast genome, and that could be cultured in standard non-selective media.

The design of our cross-species gene complementation approach required two synthetic modifications of the *POLR2I* ORF. First, the *POLR2I* ORF had to be codon-optimized to ensure proper *POLR2I* expression in *S. pombe* cells, since codon-biases differ between *S. pombe* and human [42,43]. This differs from previous *rpb9^POLR2I^* complementation approaches that used *POLR2I* derived from human complementary DNAs (cDNAs) [17,32,34], which would not have been codon-optimized for *S. pombe* expression. Second, the *POLR2I* sequence in the *rpb9*Δ*::POLR2I* strain lacked introns. While *POLR2I* has five introns in human cells [44] and the general mechanisms of RNA splicing are well-conserved between *S. pombe* and human [45,46], there are also key differences in the typical structure of introns from *S. pombe* and human [47] that could make the splicing of human introns in *S. pombe* cells not ideal. This second modification was also adopted by previous *rpb9^POLR2I^* complementation studies [17,32,34]. Since our primary goal was to endogenously express the POLR2I protein in *S. pombe*, we decided to commercially synthesize double-stranded DNA that corresponded to codon-optimized intronless *POLR2I* and integrate this version into *S. pombe* cells (Table S2). The predicted POLR2I amino acid sequence is the same between our humanized yeast and human cells.

To measure the expression and localization of POLR2I protein in our humanized *S. pombe* cells, we first tagged the C-terminus of POLR2I with green fluorescent protein (GFP) and then performed western blotting and fluorescence microscopy. As a control, we also generated *S. pombe* cells with endogenously tagged Rpb9-GFP, at the C-terminus, to allow comparison of POLR2I-GFP expression with physiological levels of Rpb9 protein expression. We find that POLR2I-GFP is expressed, although the protein level is modestly reduced compared to Rpb9-GFP (**Figure 1B**). Also, both Rpb9-GFP and POLR2I-GFP primarily localize to the *S. pombe* nucleus (**Figure 1C**), as expected for core subunits of Pol II [15]. These results indicate that *POLR2I*, which is genomically integrated at the yeast *rpb9* locus, is expressed in *S. pombe* cells to produce nuclear POLR2I protein.

### 3.2. Characterizing effects of POLR2I on S. pombe cell growth in non-selective media

Hereafter, we performed our experiments using non-selective yeast extract with glucose media (YEA), which is a standard rich media for *S. pombe* [48]. In this liquid media, we observed that *rpb9*Δ*::POLR2I S. pombe* cells grew substantially faster than *rpb9*Δ cells, but not fully as fast as wild-type (*rpb9+*) cells (**Figure 2A**). This growth pattern was also observed in the context of colony sizes where *rpb9*Δ cells often produced colonies that were smaller than wild-type ones, but *rpb9*Δ*::POLR2I* gave rise to colonies that were approximately as large as wild-type colonies (**Figure S2**). By light microscopy, we did not notice gross anomalies in the morphology of *rpb9*Δ or *rpb9*Δ*::POLR2I* cells (**Figure 2B**). These results suggest that endogenously expressed *POLR2I* can largely complement the general growth defects of *S. pombe* cells lacking *rpb9*.

**Figure 2.**
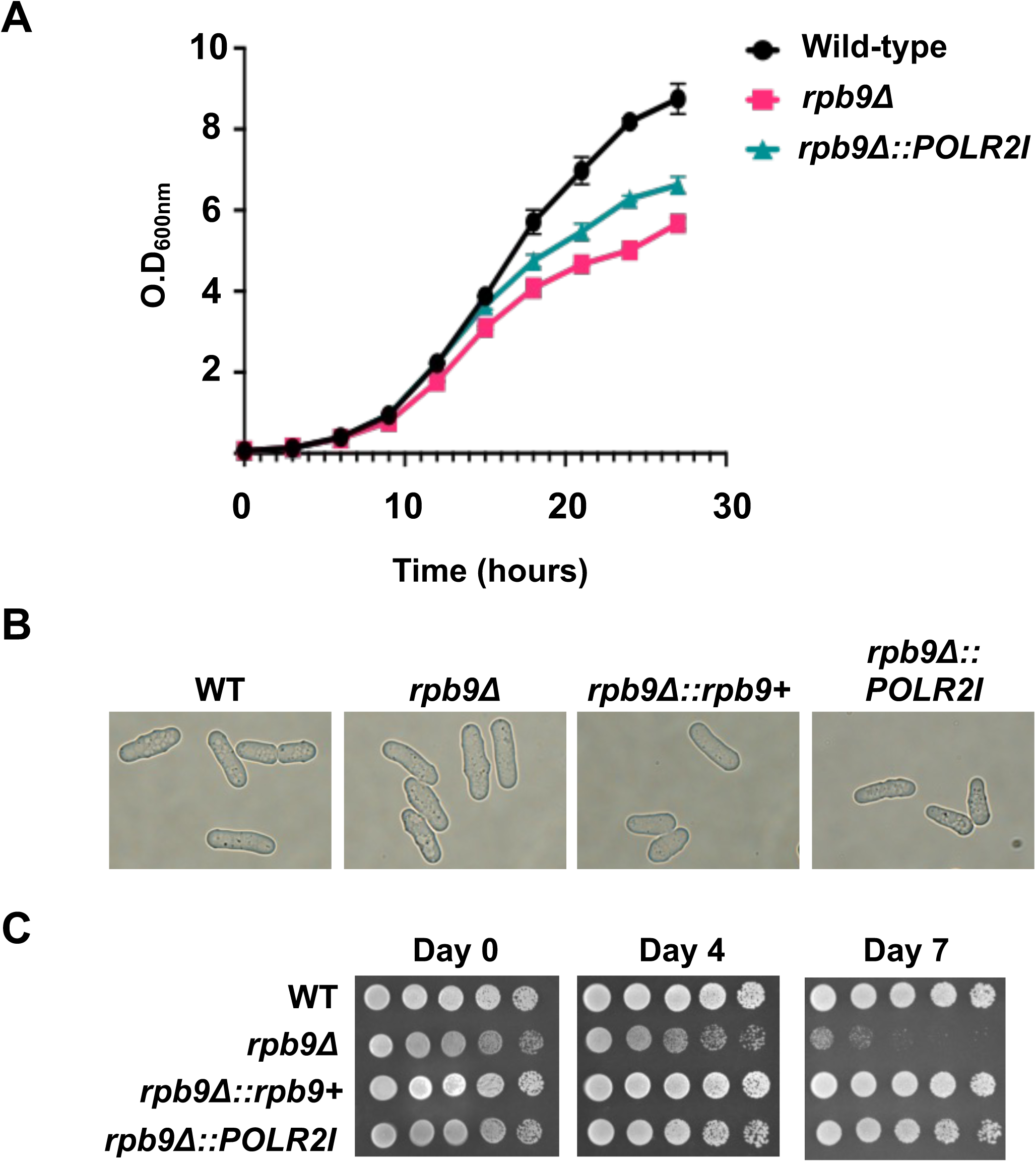
Effects of Rpb9 and POLR2I on yeast growth, physiology, and chronological aging. (**A**) Growth rate of yeast cultures as measured by optical density at 600nm wavelength readings. N = 3. (**B**) Brightfield microscopy images of cells belong to the indicated yeast strains. (**C**) Spotting assays showing the chronological aging phenotype of the indicated yeast strains, as a function of days after yeast cultures had reached stationary phase.

We also examined whether *POLR2I* could complement the function of *rpb9* in *S. pombe* aging because a recent study indicated that *rpb9*Δ reduces chronological lifespan of *S. pombe* cells [34]. Chronological lifespan is defined as the duration during which yeast cells remain viable in the stationary phase in unchanged liquid media (e.g., YEA) [49]. We tested whether *POLR2I* expression can restore the aging phenotype in *rpb9*Δ cells. We found that the viability of *rpb9*Δ cells was drastically reduced after 7 days of continuous culturing in unreplenished YEA media (**Figure 2C**). In contrast, cells where *rpb9+* or *POLR2I* was reintroduced into the native *rpb9* locus continued to grow robustly at the 7-day time- point (**Figure 2C**). These results agree with a prior study that *rpb9* loss reduces *S. pombe* chronological lifespan [34]. Furthermore, they indicate that human *POLR2I* complements the role of *S. pombe rpb9* in the chronological aging process.

### 3.3. POLR2I complements defects in stress responses and facultative heterochromatin in rpb9Δ cells

In *S. cerevisiae* and *S. pombe*, *rpb9* was found to be important for yeast viability in response to various environmental stressors [16,17]. Additionally, ectopic high expression of *POLR2I* in *rpb9*Δ cells can complement environment-dependent growth defects [17,32]. We sought to test whether endogenously expressed *POLR2I* could complement similar stress-related growth defects. We found that, while 150mM NaCl severely inhibits the growth of *rpb9*Δ cells on YEA media, growth was robustly restored for *rpb9*Δ*::POLR2I* cells (**Figure 3A**). We also tested whether *rpb9^POLR2I^* affects *S. pombe* response to the chemotherapy drug 5-fluorouracil (5-FU) because recent studies suggested that *rpb9* promotes growth of *S. pombe* cells in the presence of 5-FU [50] and that *POLR2I* gene amplification might promote 5-FU resistance in human head and neck cancers [25]. We observed that *rpb9*Δ*::POLR2I* cells could grow on YEA media in the presence of 25 μM 5-FU, unlike *rpb9*Δ cells (**Figure 3B**). Lastly, we found that endogenous *POLR2I* rescued cell growth in the presence of the antifungal drug, thiabendazole (TBZ) [51,52] (**Figure 3C**). These growth defects were also rescued by applying our humanization procedure to re-integrate *S. pombe rpb9* (*rpb9*Δ*::rpb9+*) (**Figures 3C and S3**). Altogether, these results indicate that endogenous human *POLR2I* suppresses diverse environmental-related and *rpb9*-dependent yeast growth defects.

**Figure 3.**
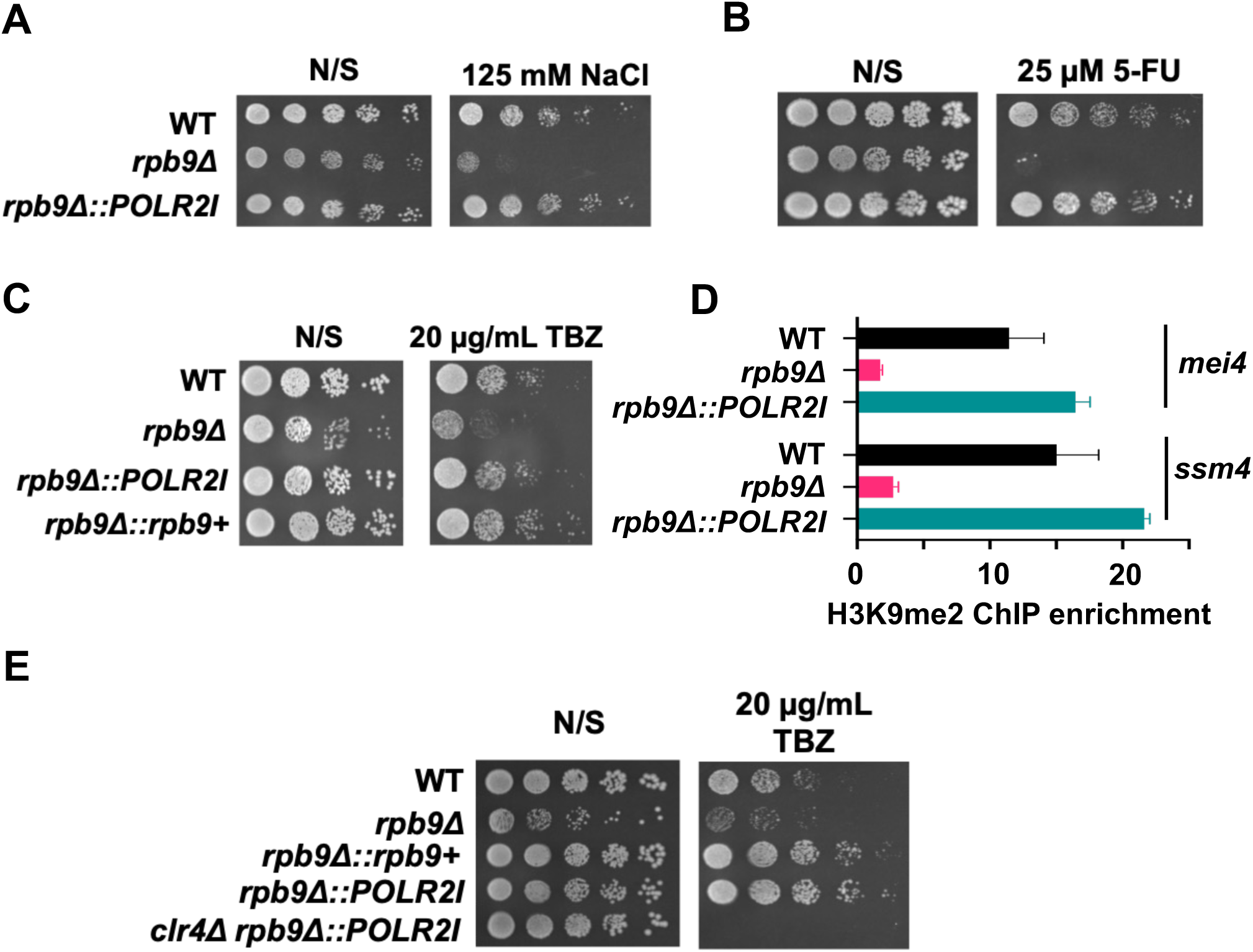
POLR2I complements rpb9 roles in environmental stresses and facultative heterochromatin formation. (**A**) Spotting assays with serial dilutions of cells on non-selective (N/S) YEA media or YEA media with 125 mM NaCl. (**B**) Spotting assays with YEA media −/+ 25μM 5-FU. (**C**) Spotting assays with YEA media −/+ 20 μg/mL TBZ. (**D**) H3K9me2 ChIP enrichments at the mei4 and ssm4 loci. Error bars denote the standard deviation of a representative ChIP experiment, N=3. (**E**) Spotting assays with YEA media −/+ 20 μg/mL TBZ. For all spotting assays, the yeast strains are indicated to the left of each panel.

Next, we explored whether yeast *rpb9* has a novel role in the formation of facultative heterochromatin in *S. pombe*, which are repressed portions of the genome that are characterized, in part, by di-methylation of histone H3 at lysine-9 (H3K9me2) [53,54]. We previously found that *S. pombe* Rpb9 can directly interact with Mmi1 [55], which is required for H3K9me2 at some facultative heterochromatin regions [54]. Therefore, we tested whether *rpb9* was required for Mmi1-dependent H3K9me2 and whether *POLR2I* can complement that putative role. By using chromatin immunoprecipitation (ChIP), we found that *rpb9*Δ abolished H3K9me2 at the *mei4* and *ssm4* loci (**Figure 3D**), which are two representative genomic regions where Mmi1 promotes H3K9 methylation [54,56,57]. Furthermore, H3K9me2 levels were restored in *rpb9*Δ*::POLR2I* cells (**Figure 3D**). This indicates that *rpb9* and *POLR2I* may share a conserved mechanism that permits the formation of facultative heterochromatin in *S. pombe* cells.

H3K9 methylation requires Clr4, which is the only H3K9 methyltransferase in *S. pombe* [58,59]. As *rpb9*Δ and *clr4*Δ cells are hypersensitive to YEA media containing TBZ drug (**Figure 3C**) [60,61], we tested whether *rpb9* mediates yeast growth response to TBZ in a Clr4-dependent manner. To do this, we deleted the *clr4* gene in *rpb9*Δ*::POLR2I* cells and assayed their growth on YEA media with TBZ. Unlike *rpb9*Δ*::POLR2I* cells that grew similar to wild-type cells (**Figure 3C**), *rpb9*Δ*::POLR2I clr4*Δ cells failed to grow (**Figure 3E**). This suggests that Clr4 likely acts downstream of yeast *rpb9^POLR2I^* to regulate TBZ responses.

### 3.4. Endogenously expressed POLR2I does not complement growth defects of rpb9Δ cells in the presence of 6-azauracil (6-AU)

Yeast Rpb9 promotes transcription elongation at transcription pause/arrest sites [19,20]. Therefore, we tested whether *rpb9* and *POLR2I* are responsive to the transcription elongation inhibitor drug 6-azauracil (6-AU), which reduces transcription elongation by depleting intracellular levels of uracil triphosphates and guanine triphosphates [62]. After knocking out *rpb9*, we found that cell growth was strongly inhibited by 6-AU (**Figure 4A**). Our finding agrees with a previous study, which showed that *rpb9* loss in the budding yeast *S. cerevisiae* reduced cell growth on media containing 6-AU [19]. Sensitivity to 6-AU was *rpb9*-dependent because reintroduction of *S. pombe rpb9* (*rpb9*Δ*::rpb9+*) led to restored yeast growth on YEA media with 6-AU (**Figure S4**). In contrast, *rpb9*Δ*::POLR2I* cells grew poorly in the presence of 6-AU (**Figure 4A**). Therefore, endogenous *POLR2I* in our humanized system is unable to compensate for the function of *S. pombe rpb9* in permitting yeast growth upon 6-AU exposure.

**Figure 4.**
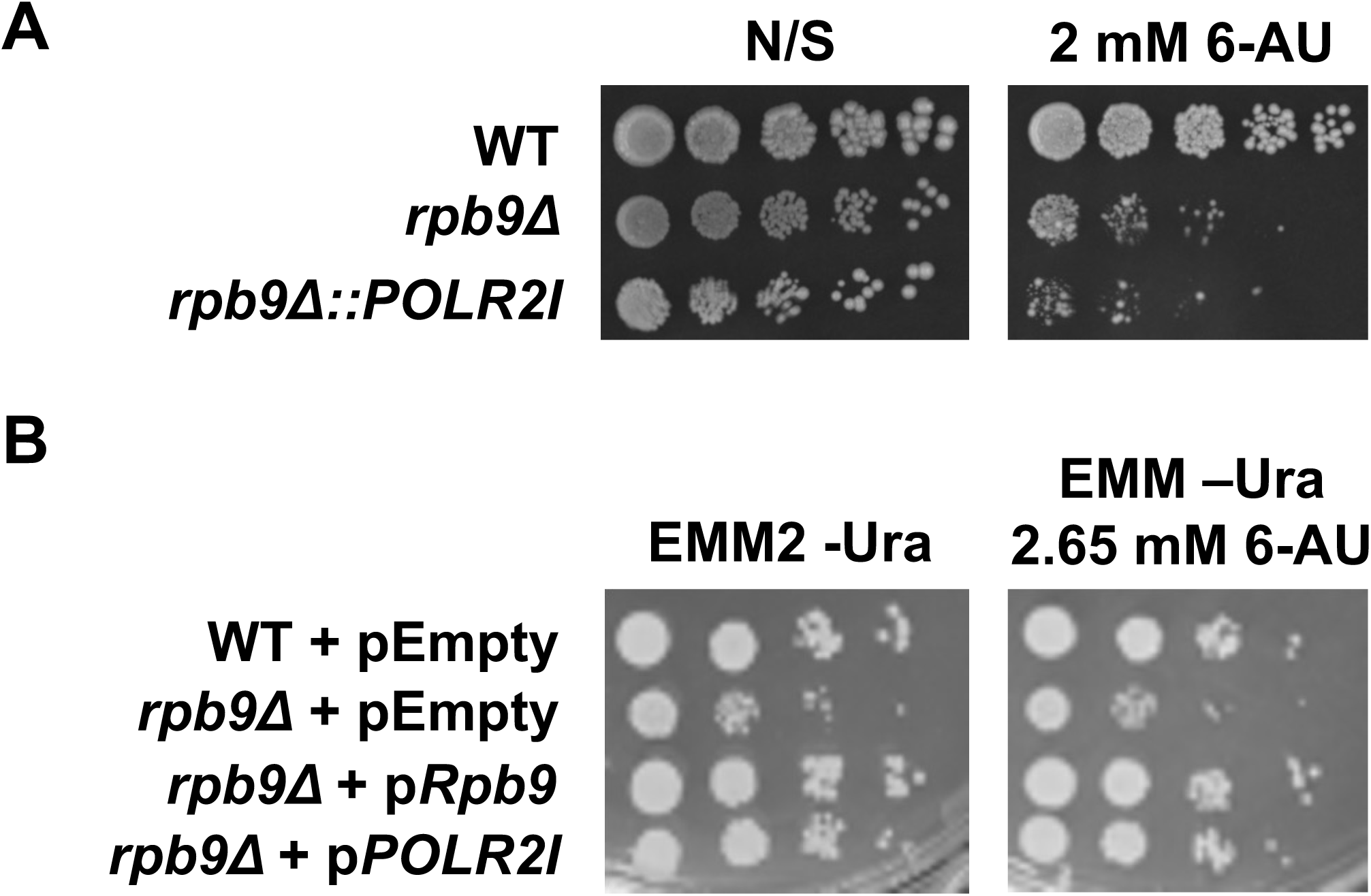
Context-dependent complementation of *POLR2I* for *rpb9*-dependent resistance against 6-AU. (**A**) Spotting assay on YEA media −/+ 2 mM 6-AU. Strains used in this panel do not carry any plasmid. (**B**) Spotting assay using cells with empty (pEmpty) or expression plasmids (p*Rpb9*, p*POLR2I).* Media lacking uracil was used to maintain selection of the plasmids, therefore ensuring that the cells retain them.

Next, we examined whether ectopic *POLR2I* could restore the growth of 6-AU-exposed *rpb9*Δ cells. Using our previously described plasmid-based approach to express *POLR2I*, from a plasmid with the P81*nmt1* promoter, in *rpb9*Δ cells [17,34], we found that ectopic *POLR2I* could complement 6-AU-associated growth defects (**Figure 4B**). This suggests that the design of cross-species gene complementation assays can impact the levels of complementation.

### 3.5. Structural similarities and divergences of S. pombe Rbp9 and human POLR2I

Protein sequence and structure are major determinants of protein functions [63]. Hence, the ability of endogenously expressed POLR2I to fully or partially complement many phenotypic roles of *S. pombe* Rpb9 suggests a high degree of conservation at the sequence and/or structural levels [17,64]. Indeed, the two homologs show 51% identity at the DNA sequence level and 47% identity at the amino acid level, based on the native sequences for *rpb9* and *POLR2I* (**Figures S5A and B**). We leveraged AlphaFold to predict structures of *S. pombe* Rpb9 and human POLR2I (**Figure 5A**). Structural comparisons of the predicted models revealed high overall similarity. Since experimentally resolved structures of Rpb9 and POLR2I already exist [12,65], we additionally compared those empirical structures (Rpb9, 3H0G-I; POLR2I, 9EHZ-F) and also found high overall resemblance (**Figure 5B**). The TM-score, a measure of protein structure similarity [66], of Rpb9 and POLR2I is 0.84 (for the Alpha models) and 0.79 (for the empirical models). These scores suggest high overall similarity of Rpb9 and POLR2I. Lastly, we used AlphaFold to predict how POLR2I could interact with the two largest subunits of Pol II, Rpb1 and Rpb2, because structural features or interactions within the larger complex may influence the functions of Rpb9 and POLR2I [55]. We focused on Rpb1 and Rpb2 because Rpb9 primarily makes direct contacts with those Pol II subunits [12]. The AlphaFold-generated models revealed notable overlaps in the predicted interactions of Rpb9 (or POLR2I) with *S. pombe* Rpb1 and Rpb2 (**Figures 5C, S5C, and S5D**). The α-carbon root mean square deviation (Cα RMSD) between the models is 0.61 Å, indicating very high similarity between the two models [67]. Upon closer inspection, two structural differences were apparent. The first was at the N-terminus of POLR2I, which has an extended tail that is absent from Rpb9 (**Figure S5C**). Second, there was a small insertion in the second zinc finger domain of POLR2I that maps to residues 104-105 of Rpb9 (**Figure S5C**). Overall, our cross-species structural analyses suggest that Rpb9 and POLR2I are highly conserved at the sequence, structure, and function levels, albeit with noticeable structural differences that could be phenotypically significant. Our conclusions based on empirical and predicted structures are overall consistent with those of prior studies that used experimentally resolved structures of Rpb9 and POLR2I [17,32,34].

**Figure 5.**
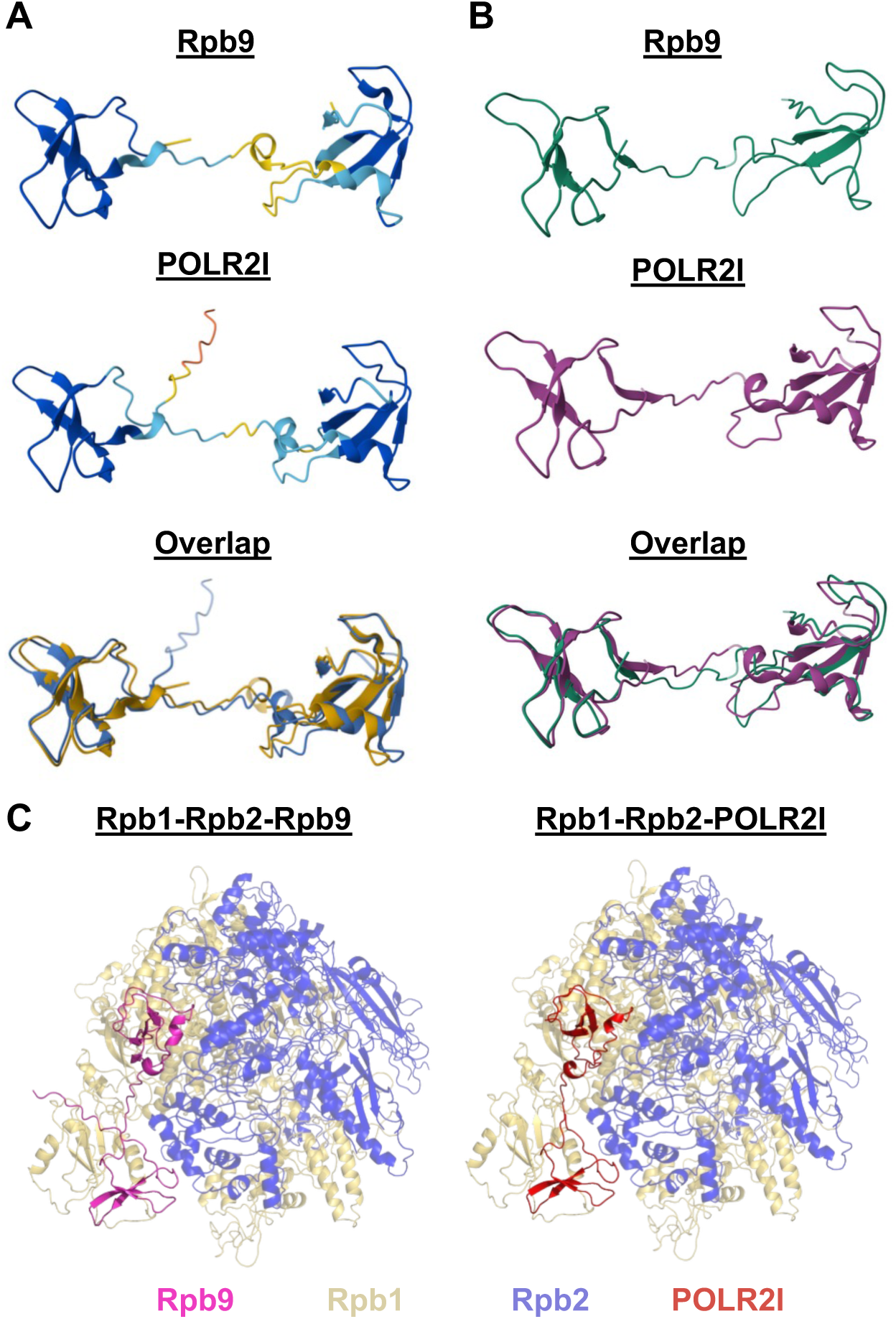
Structural analyses of Rpb9 and POLR2I. (**A**) AlphaFold-predicted structures of S. pombe Rpb9 (top) and human POLR2I (middle). Both structures are color-coded based on pIDDT confidence scores with blue (> 90): backbone and sidechain are accurate; teal (90 to 70): backbone good, side chain inaccurate; yellow (70 to 50): possible backbone inaccuracy; orange (< 50): flexible spot or intrinsically disordered. At bottom are the structures overlaid, with Rpb9 in dark yellow and POLR2I in blue. (**B**) Empirically resolved structures of S. pombe Rpb9 (top, PDB: 3H0G-I), human POLR2I (middle, PDB: 9EHZ-F), and overlaid (bottom). For the overlaid bottom structure, green is Rpb9 and magenta is POLR2I. (**C**) AlphaFold predicted structure of S. pombe Rpb9 interacting with S. pombe Rpb1 and Rpb2 (left) and AlphaFold predicted structure of human POLR2I interacting with S. pombe Rpb1 and Rpb2 (right).

## 4. Discussion

Here, we developed a novel approach to humanize *S. pombe* yeast cells to investigate the functional conservation of yeast *rpb9* and its human homolog *POLR2I*. Our approach of gene swapping directly at the genomic *rpb9* locus represents an orthogonal cross-species gene complementation strategy compared to the conventional plasmid-based approach [17,34]. We find that *S. pombe rpb9* promotes general yeast growth, growth in the presence of various environmental stressors, and histone methylation at certain facultative heterochromatin loci. While our humanization system reveals that *POLR2I* can fully or partially complement most of the defects observed in *rpb9*Δ yeast cells, *POLR2I* was unable to restore cell growth in the presence of 6-AU drug. However, this was context-dependent since plasmid-based *POLR2I* could rescue yeast sensitivity to 6-AU. Overall, our study corroborates prior conclusions from other studies that the functions of *rpb9* and *POLR2I* are highly conserved. Moreover, it reveals that the functional conservation of these two genes can vary depending on the phenotype or the assay design, suggesting context-dependency for at least some of their functions.

We observed endogenous POLR2I protein levels to be modestly lower when compared to Rpb9 levels. This was unexpected since *rpb9* and *POLR2I* were controlled by the same promoter, 5’- and 3’-UTRs, and presumably shared the same genomic context. Furthermore, the DNA sequence of the *POLR2I* ORF was codon-optimized for *S. pombe* expression. We propose that there are inherent differences in how Rpb9^POLR2I^ proteins are synthesized or stabilized in *S. pombe* cells. One possibility is that *POLR2I* is differently regulated at the transcriptional, post-transcriptional, or translational level(s). Another possibility is that the POLR2I protein is less stable compared to Rpb9, perhaps due to the absence of a human-specific stabilizing factor. These options may relate to the *POLR2I* introns that we intentionally omitted in our study design. Introns can enhance gene expression by affecting transcription, RNA transport and stability, or translation [68,69]. *S. pombe rpb9* naturally has four introns [64] and *POLR2I* may have evolved a human-specific dependence on its introns. Our unexpected finding should spur future studies into the potential role of *rpb9^POLR2I^* introns, or other factors, in regulating *POLR2I* expression. Given that *POLR2I* copy number or expression is associated with human diseases [25,26], insights into how it is regulated would be informative from a human biology standpoint. We found that *rpb9* and *POLR2I* share novel roles in H3K9 methylation at facultative heterochromatin loci *mei4* and *ssm4*. This suggests that both proteins may have additional roles in chromatin modifications, perhaps in addition to (or as a consequence of) transcription and transcription-coupled DNA repair [19,21]. Interestingly, H3K9me2 at these loci requires the *S. pombe* RNA-binding protein Mmi1 [54], which we previously found to directly interact with *S. pombe* Rpb9 [55]. Also, transcription of the *ssm4* locus is required for H3K9me2 formation at that region [54]. It is possible that *rpb9* and *POLR2I* promote Mmi1-dependent H3K9me2 indirectly through conserved transcription initiation and/or elongation functions [18,20]. Transcription would generate nascent RNAs that Mmi1 [70] could bind with to begin the H3K9 methylation process [54,56,57,71]. Alternatively, Rpb9^POLR2I^ could directly recruit or stabilize Mmi1 at the *mei4* and *ssm4* loci to promote H3K9 methylation. As *S. pombe* Mmi1 and its mouse homolog YTHDC1 [72] repress genes, in part, through H3K9 methylation [54,73], future studies into how Rpb9^POLR2I^ could be involved would improve our understanding of how RNA-binding proteins coordinate with Pol II and RNAs to regulate chromatin modifications.

Of all the phenotypes that we tested, endogenously expressed *POLR2I* failed to complement rpb9-dependent 6-AU hypersensitivity. However, reintroducing endogenous *S. pombe rpb9+* or ectopically expressed *POLR2I* restored growth, suggesting that *rpb9* and *POLR2I* might have divergent roles in their responses to 6-AU. These roles may include yeast- and human-specific mechanisms, or could relate to the small differences in protein levels within our *S. pombe* system. Alternatively, the extra N-terminal tail of POLR2I, which is absent in Rpb9, might influence 6-AU responses differently. Regardless, the mechanisms are likely distinct from those by which *rpb9^POLR2I^* regulates environmental responses or heterochromatin, in which there was robust functional complementation. Our lack of complementation finding is important because it reveals a phenotype that is distinct from the others, which is genetic support for the notion that *rpb9^POLR2I^* may be multifunctional with distinct roles [20]. Whether the role of *rpb9* in 6-AU response is generally related to transcription elongation or specifically to the 6-AU compound requires further investigation. Future studies could examine the interaction of *rpb9^POLR2I^* with genetic mutants that perturb transcription elongation.

The only other studies of functional complementation between *S. pombe rpb9* and human *POLR2I* were performed by us [17,34]. Although both our previous and current work provide largely consistent evidence for functional conservation between these homologs, we observed a discrepancy in growth responses to 6-AU. These findings underscore the value of employing orthogonal experimental approaches to dissect distinct determinants of functional conservation. While successful complementation demonstrates that homologous proteins can fulfill similar molecular roles, divergent complementation outcomes across experimental designs (such as those observed for 6-AU sensitivity) suggest that cellular and regulatory context influences how these conserved functions are executed.

## Author Contributions

Project conceptualization, T.V.V.; Designed and/or performed experiments, J.M.F., M.W., M.B.N., E.H., C.T.L., S.P. and T.V.V.; Protein structure modeling, J.M.F. and J.R.L.; Supervision, J.R.L., N.S., and T.V.V.; Wrote original draft, T.V.V.; Writing editing, J.M.F., J.R.L., C.T.L, E.H., and T.V.V.; Preparation of data figures; J.M.F., M.B.N., S.P., J.R.L, and T.V.V.; Funding acquisition, N.S. and T.V.V. All authors have read and agreed to the published version of the manuscript.

## Data Availability Statement

The yeast strains and plasmids that were used for this work are available, upon reasonable request, from the corresponding author.

## Acknowledgments

We would like to thank Matthew Faber for generating the *rpb*Δ*::kanMX* yeast strain, Dr. David Arnosti for scientific feedback on this project at group meetings, and Dr. Hyojin (Kelly) Kim for help teaching J.M.F. how to bioinformatically investigate protein structures.

---

**Figure S1.**
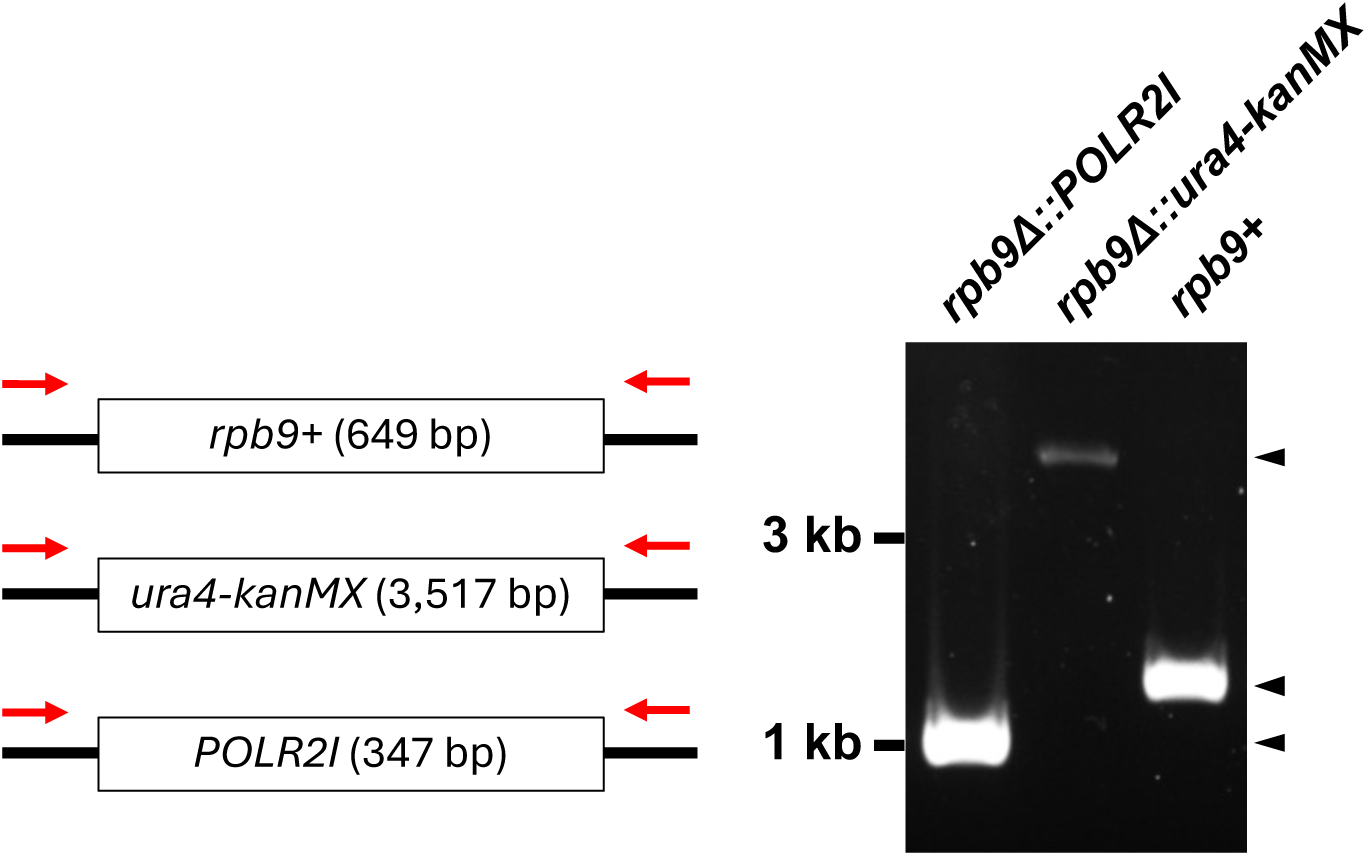
PCR-based validation of yeast strain genotypes. PCR validation of *rpb9*, *ura4-kanMX*, and *POLR2I* genes at the native *rpb9* locus in *S. pombe*. For each strain, the same set of PCR oligos was used, which targeted upstream and downstream of the swap region.

**Figure S2.**
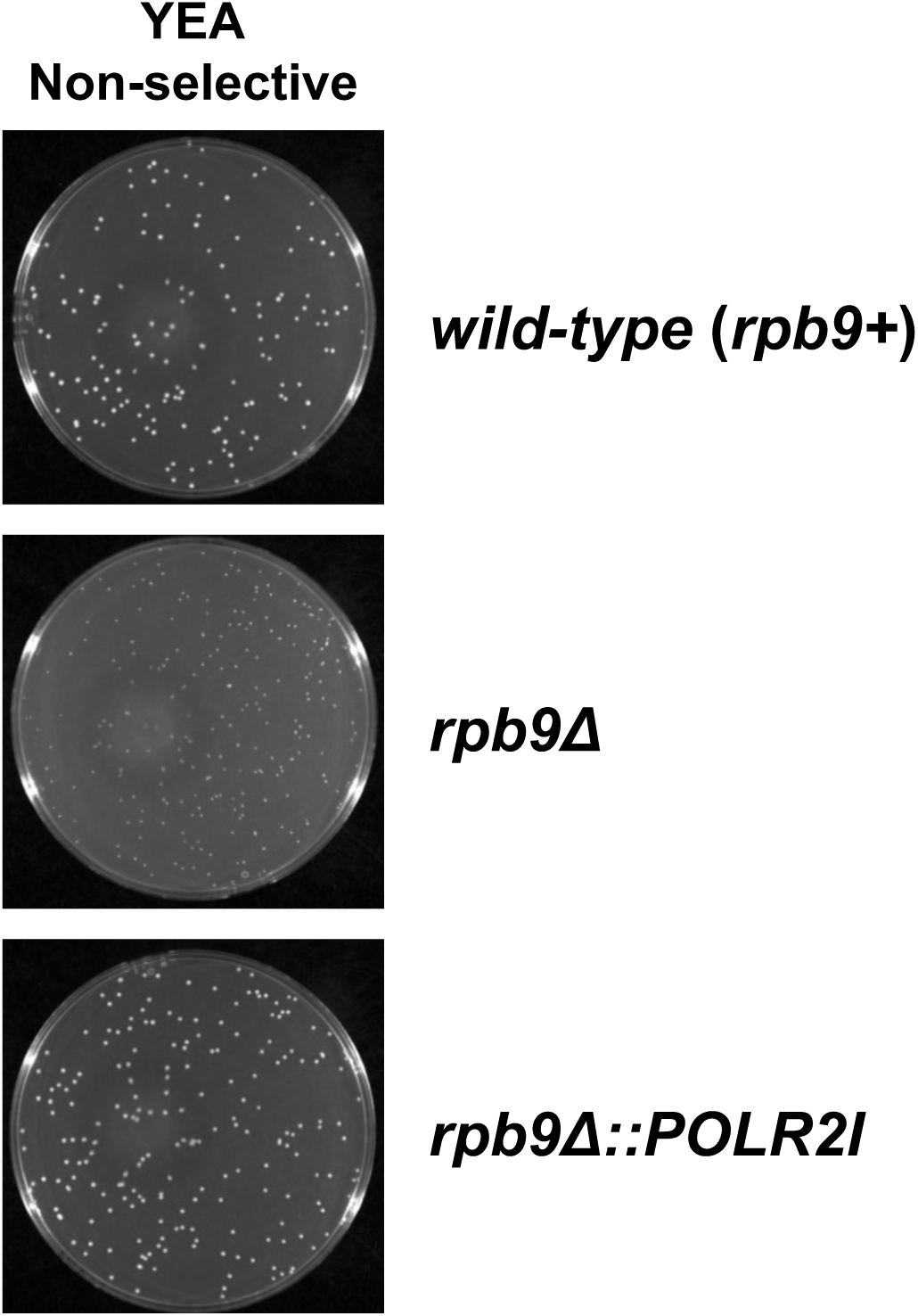
Yeast colony sizes on non-selective YEA media. Yeast colonies on non-selective YEA media plates. Approximately 1,000 yeast cells were beadspread onto the plates and incubated at 32°C for 3-5 days prior to imaging using a BioRad ChemiDoc imager.

**Figure S3.**
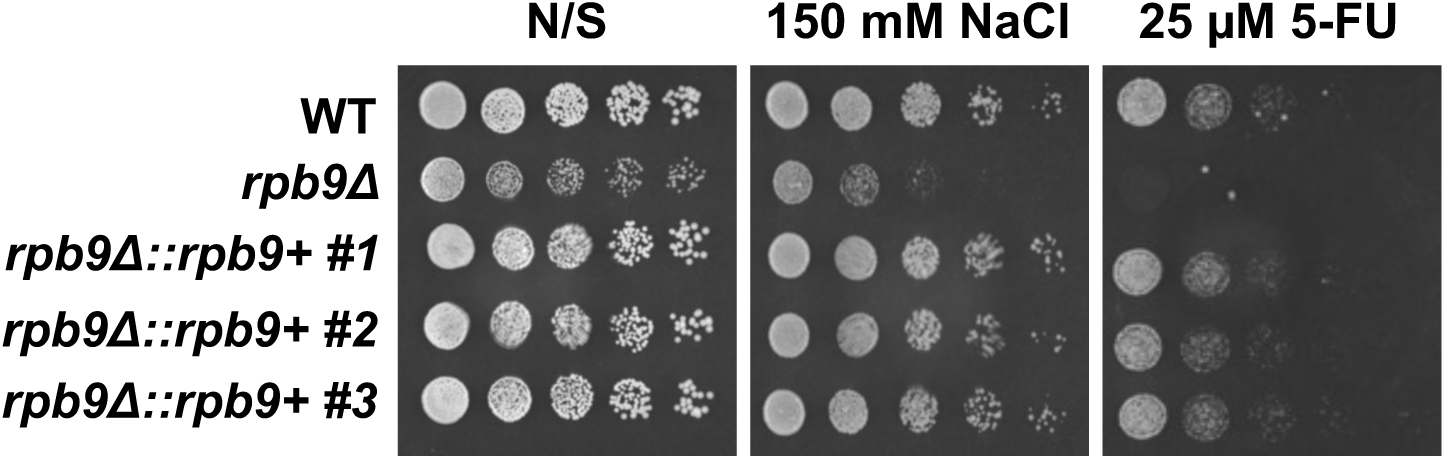
Reintroduction of *rpb9* rescues *rpb9*Δ defects. Spotting assays with YEA media −/+ 150mM NaCl or 25 µM 5-FU. Three independent isolate strains with re-integrated *rpb9+* back into the native *rpb9* locus was tested.

**Figure S4.**
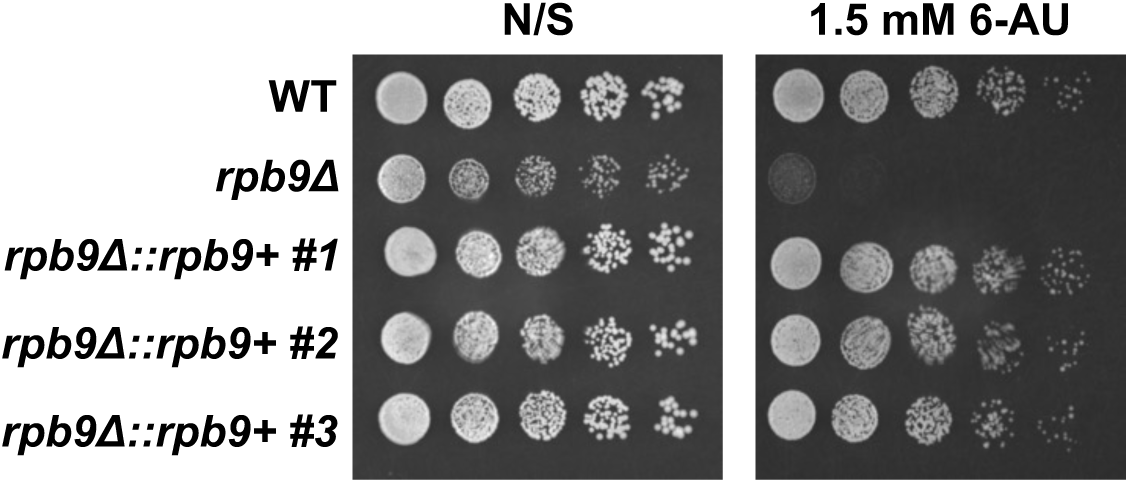
Reintroduction of *rpb9* rescues yeast growth on media with 6-AU. Spotting assays with YEA media −/+ 150mM NaCl or 1.5 mM 6-AU. The non-selective plate is identical to the one in Figure S3.

**Figure S5.**
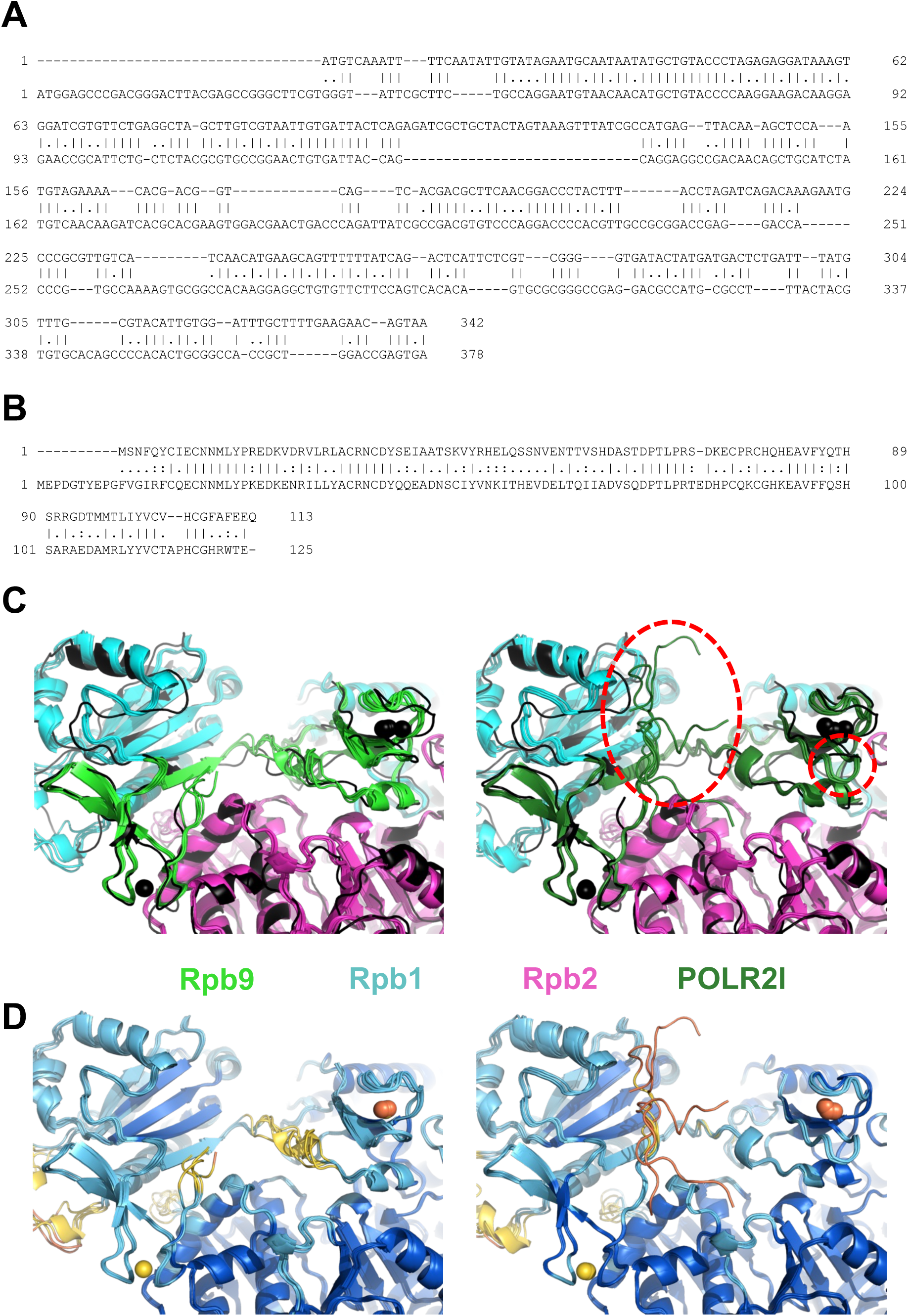
DNA and amino acid sequence alignments of *S. pombe* Rpb9 and native consensus human POLR2I. (**A**) Alignment of the DNA sequences of the open reading frames of *S. pombe rpb9* and human *POLR2I*. (**B**) Alignment of the amino acid sequences of *S. pombe* Rpb9 and human POLR2I. (**C**) (Left) The five AlphaFold3 models for the Rpb1/2/9 complex are shown with Rpb1 in cyan, Rpb2 in magenta and Rpb9 in green. The models were overlaid with the experimentally determined structure of *S. pombe* Rpb1-Rpb2-Rpb9 (black, PDB 3H0G). Black spheres indicate Zn^2+^ ions. The *C*α RMSD for the top AlphaFold3 model and 3H0G structure is 1.31, indicating very close structural similarity. (Right) Five AlphaFold3 models of Rpb1/2/POLR2I, with Rpb1 in cyan, Rpb2 in magenta, and POLR2I in dark green. Dashed red circles highlight structural differences between Rpb9 and POLR2I within those AlphaFold3 models. The N-terminus of POLR2I is likely disordered. (**D**) (Left) AlphaFold3 model of the Rpb1/2/9 complex in Fig. S5C (left) as colored based on pLDDT scores using the AlphaFold convention. (Right) AlphaFold3 model of the Rpb1/2/POLR2I complex in Fig. S5C (right) as colored based on pLDDT scores using the AlphaFold convention. The color convention is navy blue (pLDDT > 90, very high confidence), cyan (90 > pLDDT > 70, high confidence), yellow (70 > pLDDT > 50, low confidence), and orange (pLDDT < 50, very low confidence). Zn^2+^ ions are shown as spheres and are colored based on pLDDT scores.

